# Biodiversity temporal trends are reshaping food web structure and redundancy in riverine ecosystems

**DOI:** 10.1101/2025.04.23.649202

**Authors:** Alain Danet, Loubna El Madouri, Willem Bonnaffé, Andrew P. Beckerman, Maud Mouchet, Colin Fontaine, Elisa Thébault

**Author notes:** Corresponding author: Alain Danet.

## Abstract

Biodiversity temporal changes are expected to profoundly impact ecosystem functioning and stability, yet large-scale empirical evidence remains limited. Bridging this gap requires aligning biodiversity temporal trends with food web theory and Biodiversity-Ecosystem Functioning research, both of which provide strong predictions about the ecological consequences of biodiversity loss. Most insights on those consequences currently rely on spatial comparisons of ecological communities, assuming they reflect temporal trends—a space-for-time substitution. Here, we analyze biodiversity and food web time series from over 400 riverine fish communities across France (1995–2018) to evaluate how biodiversity temporal trends reshape food web structure and community functioning. We examine relationships among species richness, community biomass, and three key food web metrics: connectance, weighted average trophic level, and trophic pathway redundancy. We also test the space-for-time hypothesis using spatial gradients and a theoretical food web model. Our structural equation models and robustness analysis reveal that declines in species richness strongly correlate with reduced community biomass, with top trophic-level species playing a disproportionate role in food web changes. Declining biomass is associated with the loss of top trophic levels and decreased connectance, while temporal changes in species richness negatively correlates with changes in connectance. Both biomass and species richness declines reduce trophic pathway redundancy, suggesting that biodiversity loss weakens community robustness to future perturbations. Our findings align with food web theory and Biodiversity-Ecosystem Functioning research, support the validity of space-for-time approaches for basic food web metrics, and demonstrate the relevance of food web metrics as indicators of ecosystem function and fragility.

**Significance:** Rates of biodiversity loss raise urgent concerns about ecosystem functioning and stability, yet how biodiversity trends directly impact these critical processes remains unclear. For instance, food web & Biodiversity-Ecosystem Functioning research has delivered strong predictions about the ecological consequences of biodiversity loss but lack of empirical assessment. Using an extensive dataset of riverine fish communities, we link biodiversity trends to temporal changes in food web structure. We demonstrate that temporal changes in species richness and community biomass consistently associate with altered food web structure. We found that declining biodiversity reduces trophic pathway redundancy, likely increasing ecosystem vulnerability to perturbations. Our findings suggest that food web structure metrics could serve as valuable indicators for monitoring ecosystem health and guiding conservation strategies.

## 1 Introduction

Human pressures are driving profound biodiversity changes, including freshwater and terrestrial biodiversity declines linked to land degradation (1–3) and reductions in marine fish biomass due to intensified fishing pressures on predatory species (4). A central goal of Biodiversity-Ecosystem Functioning and food web research has been to understand the ecological consequences of these widespread biodiversity losses for ecosystem functioning and stability (5–13). However, research on biodiversity changes remains largely disconnected from studies examining their ecological impacts, primarily due to a lack of simultaneous temporal data on biodiversity and food web dynamics (14). As a result, much of our understanding of the ecological consequences of biodiversity changes over time relies on experimental and theoretical approaches (5–7, 9–11). These approaches often rely on the critical assumption that spatial comparisons of ecological communities can serve as a proxy for temporal trends—a space-for-time substitution—when direct temporal data are scarce (15). To advance our understanding of the ecological consequences of biodiversity changes, it is essential to investigate how these changes are reshaping food web structure in natural settings.

Here, we bridge the gap between biodiversity changes and food web dynamics by leveraging a unique dataset of riverine fish communities sampled repeatedly over a 23-year period (1995–2018) across 403 locations in France. By reconstructing yearly food web structures (14, 16), we address three key objectives: (i) documenting how temporal biodiversity trends reshape food web structure, (ii) connecting these changes to established theories of food webs and Biodiversity-Ecosystem functioning relationships, and (iii) assessing the validity of the space-for-time substitution assumption in the context of biodiversity and food web structure changes.

The structure of ecological networks, which describes the number, distribution, and strength of interactions among species, is critical for understanding species coexistence, ecosystem functioning, and stability to perturbations (7, 14, 17–20). Both theory and empirical evidence demonstrate that species richness and food web structure influence coexistence (17), ecosystem productivity (7, 21), temporal stability (14, 19, 20), and robustness to disturbances (9–13, 22). For instance, keystone predators can regulate dominant competitors, preventing competitive exclusion and maintaining species richness (23–25). Similarly, food webs with higher trophic levels are associated with greater ecosystem productivity (8, 21, 26, 27) and community stability (14, 19, 20), while the number and distribution of trophic links determine resilience to perturbations (9, 11, 13, 22, 28). Understanding how biodiversity changes reshape ecological networks is therefore essential for quantifying and mitigating anthropogenic impacts on ecosystems.

Three key components link biodiversity changes to food web structure. First, the energetic paradigm of food web dynamics posits that the presence of top consumers depends on sufficient resource availability at lower trophic levels, driven by bottom-up productivity (29, 30). Thus, the biomass of generalist top consumers is likely constrained by overall community biomass. Second, connectance—the proportion of realized trophic links—typically decreases with increasing species richness, as consumers are limited in the number of resources they can exploit (18, 31). However, higher species richness is expected to increase the redundancy of trophic pathways, enhancing ecosystem robustness to species extinctions (9, 11, 22, 28). This redundancy underpins the insurance effect of biodiversity, where greater diversity buffers ecosystems against perturbations and environmental changes (6, 20, 32). Finally, biodiversity-ecosystem functioning theory suggests that species richness promotes community biomass through niche complementarity (5). In food webs, the presence of higher trophic levels reduces niche overlap among producers, further boosting productivity and biomass (8, 21, 26, 27). Together, these mechanisms highlight how biodiversity changes can alter trophic link distribution, robustness, and ecosystem functioning.

To investigate how temporal biodiversity changes correlate with changes in food webs, we analyzed temporal trends in species richness, biomass, and food web structure across 6000 sampling events from 403 fish communities (Fig. 1a-c, Table S2). Using structural equation modeling (SEM), we inferred the relative effects of biomass and species richness temporal trends on trends of food web metrics, as well as the effect of species richness trends on biomass trends (Fig. 1d). Following a classical approach of network analysis (9), we also analyzed the robustness to species extinction of the meta-food web (comprising all trophic species and their interactions) to identify species extinction scenarios that could explain observed variations in food web structure. To test the space-for-time hypothesis, we compared temporal findings with spatial gradients using two approaches: (i) replicating our temporal analysis using median biodiversity and food web metrics across sites to remove temporal trends (Fig. 1c & e), and (ii) simulating food web dynamics across gradients of species richness and primary productivity using a bioenergetic model (33, 34). These simulations allowed us to evaluate whether theoretical predictions align with our empirical data, which are limited to fish biodiversity.

**Figure 1.**
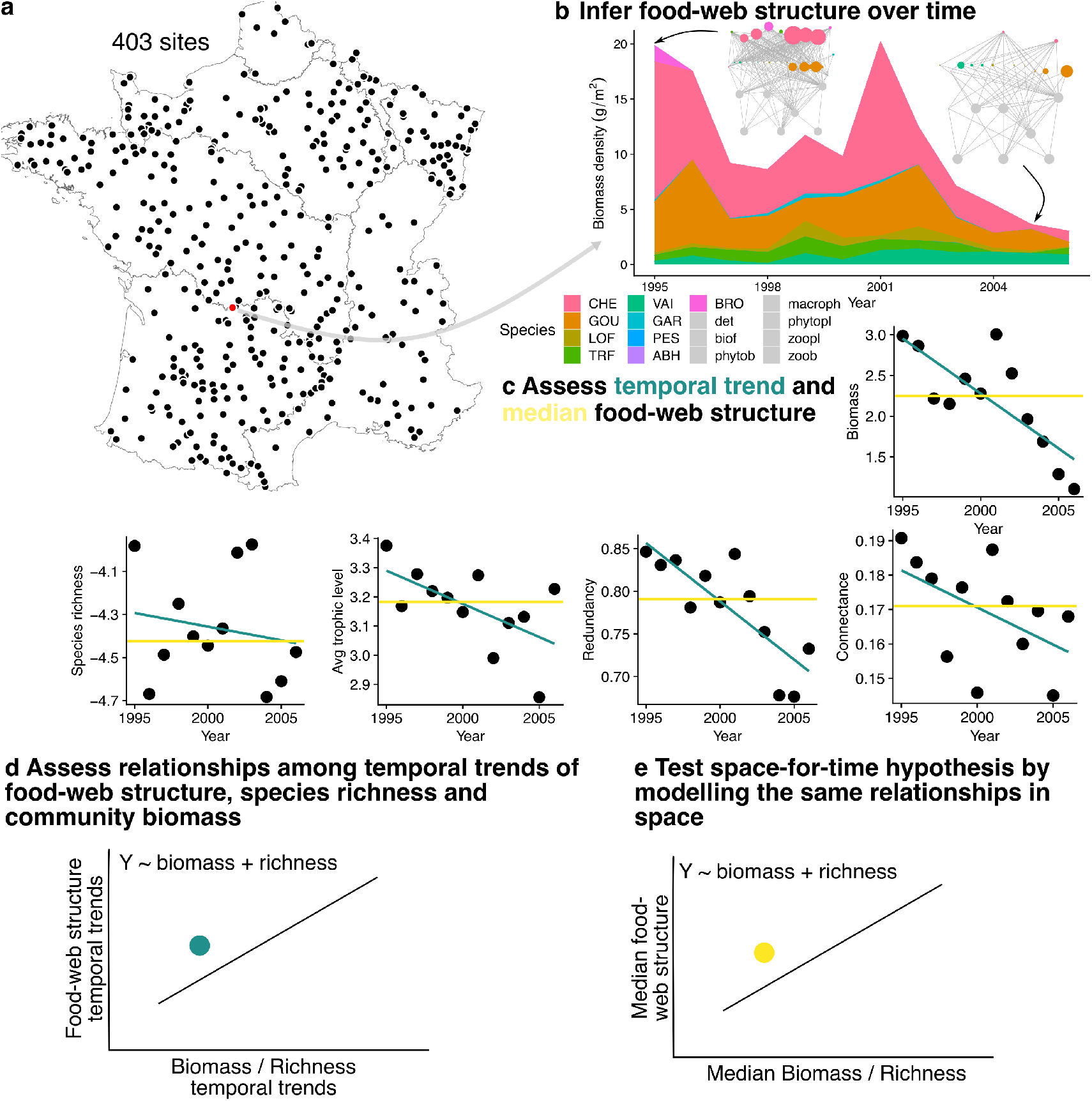
Methodological approach: (a) We analyzed 403 stream fish communities in France, each monitored for at least 10 years. (b) For each site and sampling event, we reconstructed food webs using a metaweb based on ontogenetic diet information and predator-prey body size ratios. (c) We computed temporal trends and median values for community biomass, species richness, connectance, average trophic level, and redundancy at each site. (d) We then assessed relationships among these food web descriptors over time and (e) across spatial gradients.

## 2 Results

### 2.1 Estimating temporal trends

Our food web reconstruction delivered 6000 annual estimates of species richness, biomass, connectance, weighted average trophic level, and proportion of redundant trophic links (hereafter “redundancy”, see Appendix 1, Fig. S1, Table S1) from the 403 sampling sites with 10+ years of community dynamics (Fig. 1a-c). We used ontogenic food diet data at the species level and individual fish body size to reconstruct trophic species food webs, i.e. at the size class species level (See Methods). We implemented Bayesian hierarchical models accounting for spatial variation at both hydrographic river basin and site levels to jointly estimate the temporal trends in these metrics (see Methods). We found a great heterogeneity in the temporal trends at the site level (community biomass: from -74% to +117% per decade, species richness: -52 to +64%, average trophic level: -0.31 to +0.25, connectance: -0.03 to +0.03, redundancy: -0.07 to +0.08, Fig. S2) which was effectively captured by the hierarchical model (conditional R2 ranging from 0.67 for community biomass to 0.89 for species richness, Table S3).

### 2.2 Association between biodiversity and food web temporal trends

These temporal trend estimations at the site level were the input to our SEM which accounts for uncertainties in trend estimation (see Methods). It allows us to evaluate the direct effect of the temporal trends of species richness and community biomass on the temporal trends of food web structure, the direct effect of species richness temporal trends on the temporal trends of community biomass, and the indirect effect of species richness temporal trends on the temporal trends in food web structure through community biomass. We detected a strong positive effect of the temporal trends in species richness on the temporal trends in community biomass (*r*_*δ*_ = 0.53 [0.42,0.61], Fig. 2 a and b, *r*_*δ*_: standardized slope coefficient, [95% CI], Table S4). This indicates that communities with decreasing, or increasing, species richness over time experienced a concomitant decrease, or increase, in community biomass.

**Figure 2.**
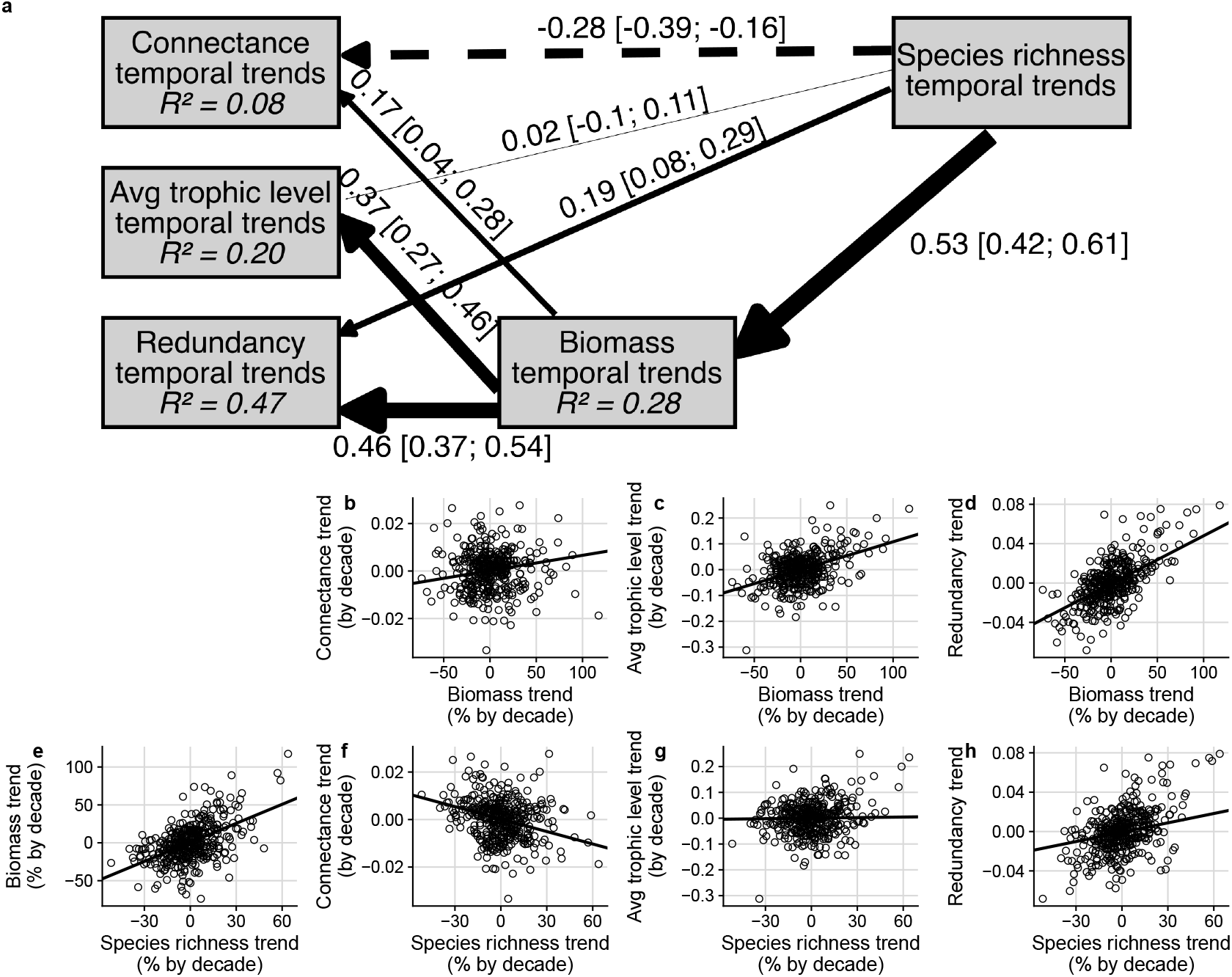
**a**, Structural equation models linking the temporal trends of biodiversity and food web structure. Dashed and solid arrows display respectively negative and positive the standardized slope coefficients, their widths being proportional to the absolute values of the coefficients. The standardized slope coefficients and their associated 95% confidence intervals obtained by bootstrapping are displayed along the path. *R*^2^ displays the marginal R-squared of the linear models. **b** to **h**, Direct bivariate relationships derived from the structural equation model. Points display the temporal trends at each site estimated with a hierarchical Bayesian model, lines display the linear relationship estimated with the SEM. The distribution of the site temporal trends are displayed in Fig. S2. All the effects are reported in Table S4. Average trophic level is weighted by the biomass of the trophic species. Redundancy is the proportion of redundant links.

We also find positive effects of the temporal trend in community biomass on the temporal trends of connectance, average trophic level, and redundancy (resp. *r*_*δ*_ = 0.17 [0.04,0.28], 0.37 [0.27,0.46], 0.46 [0.37,0.54], Fig. 2b-d). Thus, when communities experienced a decrease, or increase, in community biomass, they also experienced a concomitant decrease, or increase, in connectance, average trophic level and redundancy. To assess whether top trophic level species were linked to those relationships, we additionally looked at the temporal trends in the number and the proportion of piscivorous species. The temporal trends in community biomass were positively associated with the temporal trends in both the number and the proportion of piscivorous species (Fig. S3a). It suggests that high trophic level fishes are strongly linked to temporal trends in community biomass.

The total effect of the temporal trends in species richness on the temporal trend in connectance was negative (*r*_*δ*_ = -0.20 [-0.28,-0.10], Fig. 3a), resulting from a negative direct effect (*r*_*δ*_ = -0.28 [-0.39,-0.16], Fig. 2f) and a positive indirect effect via changes in the temporal trends in community biomass (*r*_*δ*_ = 0.09 [0.02,0.14]). Thus, when communities experienced an increase, or decrease, in species richness, they also experienced a concomitant decrease, or increase, in connectance. In contrast, the total effects of the temporal trends in species richness on the temporal trend of both trophic level and redundancy were positive (resp. *r*_*δ*_ = 0.21 [0.08,0.31], 0.43 [0.34,0.50], respectively, Fig. 3a). The total positive effect on the temporal trend of trophic level resulted from a combination of negative direct and positive indirect effect while the effect on the temporal trend of redundancy resulted from both positive direct and indirect effects (Fig. 3a). All together, this means that communities that have experienced decline in species richness over the last decades have decreased in average trophic level and redundancy but increased in connectance. Conversely, those that experienced species richness increases have increased in average trophic level, redundancy, but decreased in connectance.

**Figure 3.**
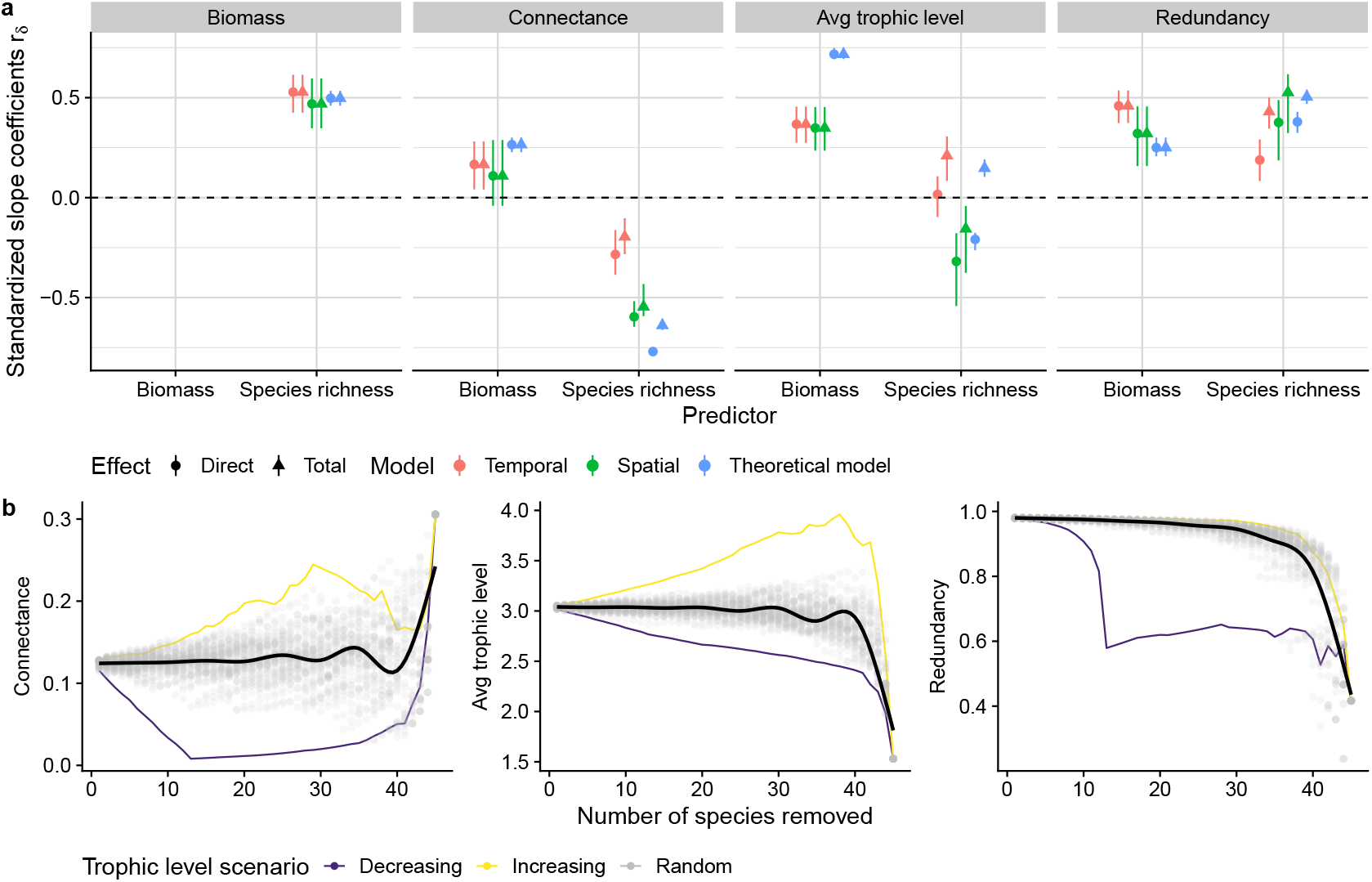
**a**, Direct and total effects of community biomass and species richness trends on food web structure trends. Standardized coefficients (*r*_*δ*_) derived from the temporal, spatial, and bioenergetic model Structural Equation Models. Points and lines display the estimated effects and their confidence intervals at 95%. Values are displayed in Table S4, *R*^2^ in Table S5, and the visualization of the bivariate relationships for the spatial and bioenergetic model SEMs are displayed in Fig. S4. Average trophic level is weighted by the biomass of the trophic species. Redundancy is the proportion of redundant links. **b**, Effect of species extinctions on the structure of the empirical metaweb according to three scenarios. In the Decreasing and Increasing scenarios, species were ordered by their trophic level and removed from the metaweb starting respectively from the highest and lowest trophic level. Species were removed in a random sequence (50 sequences, grey dots) for the random scenario, the black line displaying the average predicted by a GAM linear model. For the extinction sequence, the average trophic level was not weighted by the biomass of species as this information is not available at the metaweb scale. Similar results were obtained with trophic species (i.e. node) extinction (Fig. S5).

### 2.3 Spatial association between biodiversity and food web

Spatial analyses, either using empirical data or the bioenergetic food web model, largely aligned with temporal trends (Fig. 3a and S4), showing consistent positive effects of species richness on community biomass and of community biomass on average trophic level and redundancy. Across space, sites with higher fish biomass on average had food webs with higher connectance, redundancy and average trophic level than sites with lower fish biomass. The theoretical model predicted similar relations, with a particularly strong positive association between total biomass and average trophic level across modelled food webs. Species richness also exhibited a direct negative effect on connectance and a positive effect on redundancy, mirroring temporal patterns. Both modelled food webs and local fish communities characterized by greater species richness on average showed higher average biomass and food web redundancy, and lower connectance. However, spatial and temporal analyses diverged in their effects of species richness on average trophic level, with spatial data showing a negative effect and temporal trends indicating a positive one. Detailed statistical effects and comparisons are provided in the Appendix.

### 2.4 Robustness of food web structure to species extinctions

To investigate whether top species were drivers of biomass and structure patterns (see section 4.2), we implemented three secondary extinction simulation scenarios on the empirical metaweb defined as a food web containing all the trophic species, i.e. the fish species subdivided in size classes added to their resources, found across all sites in the empirical data (See Methods). When trophic species were removed in a sequential order beginning by the highest trophic levels, connectance, average trophic level, and redundancy decreased (purple lines, Fig. 3b). On the other hand, removing low level trophic species first increases connectance and average trophic level, but does not affect redundancy (yellow lines, Fig. 3b). In turn, species extinction at random had little effect on food web structure (black line, Fig. 3b). These results, in conjunction with our empirical analyses above, suggest that top consumers play a disproportionate role in driving the changes in food web structure reported.

## 3 Discussion

Our findings reveal several key patterns: (1) temporal trends in species richness were positively associated with trends in community biomass; (2) temporal trends in community biomass were positively linked to trends in connectance and average trophic level; (3) temporal trends in species richness were negatively associated with trends in connectance and only weakly related to trends in average trophic level; and (4) both community biomass and species richness trends were positively associated with trends in trophic pathway redundancy. Additionally, we found evidence that (5) top predator species play a significant role in driving these patterns. Importantly, these temporal relationships were largely consistent with patterns observed across spatial gradients, both in empirical data and in simulations from a bioenergetic food web model. In the following, we discuss these findings in the context of food web assembly theory and Biodiversity-Ecosystem Functioning theory, and consider their broader ecological implications.

We found that community biomass scaled positively with species richness, a pattern consistent across our temporal, spatial, and theoretical analyses. This result aligns with previous studies on fish communities across space (14, 35) as well as grasslands, salt marshes, and forests (36–39, but see 40). These empirical findings are supported by theory, suggesting that food web diversity enhances biomass by stimulating primary production and resource uptake (8, 21, 26, 27), and by Biodiversity-Ecosystem Functioning theory, which predicts that species loss reduces community biomass (5). Our results bring a temporal dimension to these findings using long-term trends in empirical data, and clearly exemplify that declines in species richness goes along with a reduction of biomass in ecosystems. Conversely, our result also indicates that conservation actions promoting species richness should also promote community biomass.

Our results also reveal that food webs experiencing biomass and/or species richness declines undergo significant structural changes, including the loss of top predators and a reduction of connectance. These findings align with energetic and trophic biogeography theories (29, 30), which posit that the presence of high-trophic-level species is constrained by the availability of energy and resources (41). Thus, a decline in community biomass may lead to insufficient resources to sustain top predators, while an increase in biomass could provide the necessary energy to support these species (41, 42). Further, Biodiversity-Ecosystem Functioning theory suggests that the presence of higher trophic levels may reduce niche overlap among producers, enhancing productivity and community biomass (8, 21, 26, 27). The relationship between community biomass and connectance is less clear, with a weaker positive association compared to average trophic level. Previous theoretical studies have reported contrasting effects of connectance on productivity (7, 8, 26, 27, 43). For instance, while increased connectance between herbivores and plants can either enhance or reduce productivity (7, 27, 43), greater connectance among carnivores generally boosts productivity by reducing top-down control and allowing biomass to accumulate across trophic levels (8, 26). Interestingly, our theoretical model predicts overall a positive relation between total biomass and connectance, which contrasts with the negative relation found recently by another complex allometric food web model (27), but this relation is far weaker than the strong positive relation between total biomass and average trophic level.

Our study suggests that declines in community biomass and species richness are associated with a loss of redundant trophic links, while their increase results in greater redundancy. Since redundant links are those that can be removed without triggering secondary extinctions (22, 28), our findings suggest that declines in community biomass and species richness might reduce food web robustness to further perturbations (11). Such positive effect of species richness on redundancy would then further support the insurance hypothesis (32, 44), where greater species richness and functional redundancy buffer ecosystems against perturbations, increasing stability (13). Furthermore, our results highlight the disproportionate role of top trophic-level species in maintaining redundant links within food webs. This pattern is consistent with previous empirical studies showing that top predators are often the most generalist and highly connected species (12, 45). Consequently, disturbances affecting these top predators are likely to have the greatest impact on overall community stability (12). These findings underscore the importance of preserving top predators and maintaining species richness to safeguard food web robustness and ecosystem stability.

Species richness was negatively associated with connectance, which is consistent with previous findings and likely driven by limitations in predator generality (31). In contrast, the effects of species richness on average trophic level were more nuanced and considerably weaker than those of community biomass. Direct effects of species richness on average trophic level were either negligible or negative, as observed in temporal, spatial, and theoretical models. However, when accounting for the indirect effects of species richness — mediated through changes in community biomass — our temporal and theoretical analyses revealed a positive association between species richness and average trophic level. This suggests that while species richness influences food web structure, its effects are more pronounced on connectance than on average trophic level dynamics.

We found strong agreement between our temporal and spatial analyses, as well as with predictions from food web theory and Biodiversity-Ecosystem Functioning research. This suggests that spatial comparisons of generic topological food web properties and biodiversity metrics may serve as reasonable surrogates for predicting temporal trends—supporting the validity of the space-for-time substitution in both empirical data and bioenergetic food web models. However, this agreement did not hold for the relationship between weighted average trophic level (hereafter “average trophic level”) and species richness. This discrepancy may arise because average trophic level incorporates both food web topology and biomass distribution, which are influenced by dynamic processes such as bottom-up and top-down control. Indeed, biomass distribution across trophic levels depends on species metabolism and productivity which in turn control the strength of top-down and bottom-up processes (33, 46).

Our study provides compelling evidence that biodiversity trends are profoundly reshaping food web structure. By linking these patterns to Biodiversity-Ecosystem Functioning and food web theory, we demonstrate that widespread biodiversity declines likely have significant and detrimental consequences for ecosystem functioning and stability in the face of future perturbations. These findings suggest that assessments of food web structure—based on metaweb reconstruction like in the present study—, added to more classic metrics of community biomass, and species richness could serve as valuable indicators of functional decline and metrics for evaluating the efficacy of mitigation strategies. Finally, our study also highlights critical avenues for future research. For instance, integrating food web metrics into conservation planning will require understanding how they relate to other facets of biodiversity, such as the presence of non-native species, and drivers of biodiversity changes, including habitat degradation (3) and climate change (47).

## 4 Material & Methods

We used a monitoring of freshwater fishes conducted all over France by the Office Français de la Biodiversité (OFB). The sampling protocols were standardized since 1995 and a complete description of the sampling process can be found in (14). The fishes are sampled using electrofishing all across the streams in shallow streams and over the bank by boat in deeper streams. All the fishes are counted and the body size of fishes are recorded on a sample of individuals, enabling to estimate the distribution of individual body size by species for each sampling (14, 16).

We selected the sites that have been monitored for at least 10 years in the 1995-2018 period, with the same sampling protocol over time. We selected one sampling per year by site. In each site, we ensured low temporal heterogeneity within the sampling period by selecting samplings that were performed in the same period, leading to a mean standard deviation of sampled month in each site of three weeks (14). Those steps lead to the selection of 403 sites (Fig. 1a, Table S2).

### 4.1 Biomass

We first estimated the body mass of each fish individual with the allometric law: 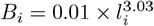, *B*_*i*_ being the biomass expressed in gram and li the body length of the individual *i* expressed in millimeters (14, 48). Then the biomass of each species and community were obtained by summing the body mass of individuals. This resulted in a biomass score for the whole community of each site. Finally, the biomass was reported to the sampled surface (in *m*^2^) to account for sampling effort.

### 4.2 Food web reconstructions

We reconstructed food web structure in two steps (14, 16): (1) we first computed the metaweb, describing the potential trophic interactions among all size classes species, (2) then we extracted yearly species size classes and their interactions according to their presence in a given sampling event, to establish a annual snapshot of food webs.

We built the metaweb according to ontogenic diet data and body size (14, 16, 43, 47, 49, 50). The method is extensively described in (16) and (14). The food diet of stream fishes changes along their life cycle or stage, therefore according to their body size. Each fish species was divided in nine size classes, i.e. trophic species, equally distributed in their observed body size range. Thus, a trophic species is a group of individuals of the same species and size class that are expected to share the same prey and predators because trophic interactions are largely determined by predator-prey size ratio (33, 50). We also added seven nodes present in the ontogenic food diet database: detritus, biofilm, phytoplankton, zooplankton, macrophages, phytobenthos and zoobenthos. The resulting metaweb contained 45 fish species divided into 9 size classes plus 7 resource nodes, summing up to 412 nodes. Consideration of size classes, i.e. trophic species instead of species, improves inference of trophic interactions (51).

We determined the feeding interactions of trophic species according to their species identity and their body size. The diet database described two or three life stages by fish species, each one being delimited by body size. The ontogenic fish diet database was filled according to fishbase (52) and literature (14, 16).

The feeding interactions among fishes were based on the body size ratio between predators and preys. We defined the predation window of a piscivorous trophic species as 3% to 45% of its midpoint (53–55). A trophic link was defined between the piscivorous trophic species and all the trophic species whose midpoint in body size range was included in the predation window. The fish-resource trophic links were set between the trophic species and the resource nodes according to the food items of their size class. Finally, the feeding interactions between resources were defined according to the literature (55, 56).

### 4.3 Community and food web structure

At each site, yearly food webs were described by their community biomass and species richness. Yearly food web structure was described by their connectance, describing the degree of generalism in a network (31, 57). Connectance was computed as the ratio between the number of actual links and the number of potential links 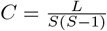, with *L* being the number of links and *S* the number of trophic species). We also computed the average trophic level, weighted by the biomass of the trophic species. The trophic level of a trophic species was computed recursively as 1 added to the mean trophic level of the preys it consumes (58). We computed the number of redundant and functional links using the dominator tree approach (11). A species *i* is a dominator of species *j* if species *i* mediates the connection of species *j* to basal resources such that the extinction of i removes the connection of *j* to basal resources. It is then used to identify the set of trophic links that are essential to connect species to basal resources, i.e. functional links, and the redundant links. This method requires a single basal root in the food webs. To do so, we added a basal dummy resource node to the empirical food webs connected to biofilm and phytoplankton resource nodes. For food webs generated from theoretical simulations (see below), we added a resource node connected to all primary producers. We computed dominator trees using the `dominator_tree()` function from the `igraph` R package (See Appendix for details). We computed the number of functional links as the number of links in the resulting dominator tree of the food web. We checked that the procedure reproduced the examples presented in previous studies (11, Appendix). We then derived the proportion of redundant links in food webs of empirical and theoretical simulations.

### 4.4 Temporal trends of food web

We modeled the temporal trends of food web structure at each site (*Y*_*i*_) using a linear hierarchical model which accounts for the hierarchical spatial structure of the data, i.e. the sites nested in hydrographic basins. Time was considered as the number of years since the first sampling (*t* = 0) at each site. *α* is the intercept and *β* the slope estimating the temporal trends. The intercept and the slope parameters have a fixed (*α*_0_, *μ*) part and a random part. We accounted for the spatial structure of the data by adding random intercepts (*a*) and slopes (*b*) for the basin (*a*_*n*_, *b*_*n*_) and site (*a*_*i*|*n*_, *b*_*i*|*n*_) identity, site effects being nested in the basin one.

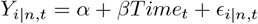

- *α* = *α*_0_ + *a*_*n*_ + *a*_*i*|*n*_
- *β* = *μ* + *b*_*n*_ + *b*_*i*|*n* 2_
- *a*_*n*_, *a*_*i n*_, *b*_*n*_, *b*_*i* | *n*_, *ϵ*_*i* | *n*,*t*_ ∼ 𝒩 (0, *σ*)
- *n*: basin, *i*: site *i, t*: time *t*

The model parameters were estimated using INLA (Integrated Nested Laplacian Approximation) Bayesian inference with default uninformative priors (59). The prior distributions for the fixed intercept (*α*_0_) and slope (*μ*) were modeled as a flat normal distribution with a mean of zero (𝒩 (0, 1000)). Prior distributions for the Gaussian error *ϵ*_*i*|*n*,*t*_ and the random effects were modeled as a log gamma distribution with shape and inverse scale parameters (𝒢 (*s, τ*) = 𝒢 (1, 5.10^*−*^5)), then their respective standard deviation were obtained by back-transforming the inverse scale parameter 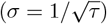. Community biomass and species richness were log-transformed.

The performance of the model was assessed visually by plotting the fitted versus the observed values (Fig. S6). We also computed the marginal 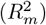 and conditional 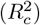 R-squared of the models following previous formulations (60, 61). In order to account for the variability associated with the priors (62), we computed the variance of the posterior distribution of each predicted values (*V ar*_*fit*_ = *σ*^2^(*Ŷ*_*i*_)). We computed *R*^2^ values for each predicted values, then obtaining a *R*^2^ distribution (62). We reported mean marginal and conditional *R*^2^ associated the 95% credible interval computed using the Highest Posterior Density method.

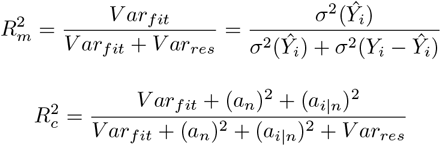

*y*_*i*_ and *ŷ* _*i*_ being respectively the observations and the predicted values, *V ar*_*fit*_ and *V ar*_*res*_ being respectively the variance of predictive means and the variance of the residuals. *a*_*n*_ and *a*_*i n*_ are respectively the standard deviation on the random intercept associated to the basin and the site.

We extracted the annual temporal trends at each site from the hierarchical model using mean Best Linear Unbiased Prediction (BLUP) values, then multiplied by 10 to get temporal trends by decade. For community biomass and species richness which were log-transformed, we derived temporal trends in percentage change by decade by back-transforming the slopes (*β*′ = (*e*^*β*^ − 1) × 100 × 10).

Overall, we found that standard deviation attributed to site effects in both baseline (i.e. the intercept) and temporal trends in community structure were much higher than the ones found for hydrographic basin (Table S6).

### 4.5 Spatial

We assessed the validity of the space-for-time substitution hypothesis by comparing the relationships estimated among food web structure in time to the ones inferred in space. We used the same sites as in the temporal analysis, but we summarized the temporal series of each station by the median value over time of each facet of food web structure described above (Fig. 1).

### 4.6 Statistical analysis

We modeled the relationships among food web structure indices, i.e. connectance, average trophic level, the proportion of redundant trophic links, and community biomass and species richness, with Structural Equation Models. We modeled species richness as related to community biomass and food web structure. In turn, community biomass was related to food web structure. We computed one structural equation model to assess the relationships among the temporal trends and one for the relationships over the spatial gradient, we added a random effect of the hydrographic basin on the intercept for the latter one.

In order to control for the uncertainties of the estimated temporal trends, we weighted each temporal trend by the inverse of the sum of the standard deviation of the estimated trends, i.e. the estimated slopes for the response and the explicative variables (*w*_*j*_ = 1/ ∑*s*_*j*_, *w*_*j*_ being the weight, and *s*_*j*_ the standard deviation on the temporal trends of the variables).

The coefficients of the Structural Equation Models were computed with PiecewiseSEM R package (63). The coefficients were standardized and the confidence intervals were computed with semEff R package (64). Confidence intervals were obtained by bootstrapping the linear models 1000 times and computing adjusted bootstrap percentile intervals (BCa), which are adjusted for the asymmetric distribution of resampling. The marginal and conditional *R*^2^ associated with the linear models of the SEMs were computed with the Nagelkerke method (63, Table S5).

All the statistical models displayed low multicollinearity (Variance Inflation Factors < 3, Table S 7). We visually inspected the residuals of the linear models.

### 4.7 Extinction simulation in the metaweb

We simulated extinction sequences in the metaweb according to three scenarios to get expectations on how the food web structure should change according to the addition and deletion of species (results presented in Fig. 3b) and trophic species (results in Appendix), i.e. species subdivided in size classes. The scenario of extinction sequences were related to the trophic level of the species/trophic species. In the decreasing and increasing scenario, we removed species/nodes in decreasing, resp. increasing order, of their trophic level, i.e. respectively beginning by the highest and lowest trophic level. In the random extinction sequence, we built 50 random extinction sequences. For each simulation, we removed fish species/nodes t hat lost all their prey. We then computed the connectance, the average and the maximum trophic level after each extinction.

### 4.8 Theoretical simulations

We used a bioenergetic food web model to simulate the relations between community biomass, species richness and food web structure. This food web model lifts one of the limitations of our data, that is it includes the biomass of all species in the food webs from primary producers to top predators. This model allows simulating complex communities because its parametrization depends mainly on species body masses. We first generated 176 food webs varying in species richness (range: 10-100) a nd connectance (range: 0.02-0.32) using the niche model algorithm, which reproduces food web topology typically found in empirical food webs (65). We set species body masses (*M*_*i*_) using a predator-prey mass ratio of 100 (*Z*) (33, 50), such as 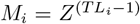, *T L*_*i*_ being the trophic level of the species *i*. The metabolic rates of the species (*x*_*i*_) were then derived from allometric relationships between metabolism and body mass empirically estimated (33), such as 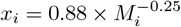.

The dynamics of species biomasses are described by an ODE systems of two equations, for primary producers (eq. (1)) and consumers (eq. (3)) respectively:

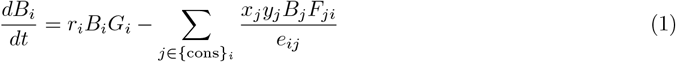

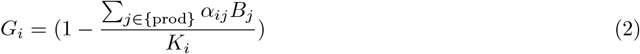

The primary producer biomass increases with a growth rate (*r*_*i*_) of 1 and a logistic growth (*G*_*i*_, eq. (2)), growth being limited by intraspecific (*α*_*ii*_, set to 1), interspecific (*α*_*ij*_, set to 0.5) competition, and carrying capacity (*K*_*i*_, varying in the simulations). Primary producers loose biomass by being eaten by consumers at a rate that depend on consumer metabolic rates (*x*_*j*_), attack rates (*y*_*j*_, set to 4 (34)), a functional response (*F*_*ji*_), and assimilation efficiency accounting for the fact that consumers assimilate better the primary producer biomass than the consumer ones (*e*_*ij*_, set to 0.45 for herbivores and 0.85 for carnivores, (34)).

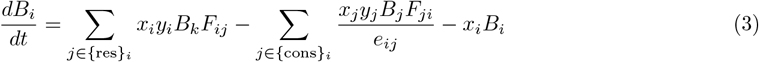

Consumers gain biomass by eating other species and lose biomass by being consumed and by metabolic loses (*x*_*i*_, eq. (3)). The functional response (eq. (4)) of consumer feeding depends on the preference for its preys (*ω*_*ij*_, set even across consumer preys) and half-saturation rate (*B*_0_, set to 0.5). The hill coefficient (*h*, set to 1.5) determines the shape of the functional response (Holling’s type II or III).

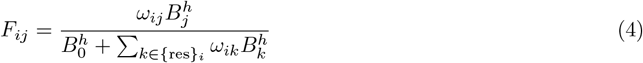

We simulated the dynamics of the 176 food webs across a gradient of global carrying capacity (*K*, range: 1-10) to represent variation in space of productivity. The carrying capacity were standardized according to the number of producers and interspecific competition 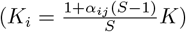to prevent changes in productivity due to system size and complementarity among producers (66, 67). We ran simulations for 5000 timesteps. At the end of simulations, we removed dead species (biomass < 10^-6) and measured community biomass, species richness and food web structure. If the final community contained disconnected producers (i.e. without consumers) or disconnected consumers (i.e. consumers without preys), we discarded them and run the simulations again with the same parametrization. The simulations were done using the Julia package `EcologicalNetworksDynamics.jl` (34). Then, we computed the community and food web metrics at the last timesteps and analyzed the data with the same SEM described above.

## 5 Data, Materials, and Software Availability

The raw data used for the analysis are available on Zenodo: https://zenodo.org/records/5095656. The raw fish database is public and available on Zenodo: https://zenodo.org/records/8099409. The manuscript and the supplementary materials are written in Rmarkdown, i.e. combining code and text, and are available on github: https://github.com/alaindanet/fishcom_biomass_dynamics. We further implemented a code pipeline using the targets R package to ensure that the code, the data, the figures, the manuscript and the results are up to date.

## 6 Acknowledgments

We are indebted to the OFB for providing the database, and we thank the numerous people who contributed to the fish records. We thank Dr Katherine Assersohn for her constructive feedback on the study. AD, LEM, MM, CF, and EL have received support from by the ANR project ECOSTAB (ANR-17-CE32-0002) awarded to EL. AD and APB have received support from the Natural Environment Research Council under grant agreement no. NE/T003502/1 awarded to APB.

## References

1. T. Newbold, et al., Global effects of land use on local terrestrial biodiversity. Nature 520, 45–50 (2015).

2. B. J. Cardinale, A. Gonzalez, G. R. H. Allington, M. Loreau, Is local biodiversity declining or not? A summary of the debate over analysis of species richness time trends. Biological Conservation 219, 175–183 (2018).

3. A. Danet, X. Giam, J. D. Olden, L. Comte, Past and recent anthropogenic pressures drive rapid changes in riverine fish communities. Nature Ecology & Evolution, 1–12 (2024).

4. V. Christensen, et al., A century of fish biomass decline in the ocean. Marine Ecology Progress Series 512, 155–166 (2014).

5. M. Loreau, et al., Biodiversity and ecosystem functioning: Current knowledge and future challenges. science 294, 804–808 (2001).

6. M. Loreau, A. Hector, F. Isbell, The Ecological and Societal Consequences of Biodiversity Loss, 1st Ed. (Wiley, 2022) 10.1002/9781119902911 (July 11, 2024).

7. É. Thébault, M. Loreau, Food-web constraints on biodiversity-ecosystem functioning relationships. Proceedings of the National Academy of Sciences 100, 14949–14954 (2003).

8. F. D. Schneider, U. Brose, B. C. Rall, C. Guill, Animal diversity and ecosystem functioning in dynamic food webs. Nature Communications 7, 12718 (2016).

9. J. A. Dunne, R. J. Williams, N. D. Martinez, Network structure and biodiversity loss in food webs: Robustness increases with connectance. Ecology Letters 5, 558–567 (2002).

10. R. V. Solé, M. Montoya, Complexity and fragility in ecological networks. Proceedings of the Royal Society of London. Series B: Biological Sciences 268, 2039–2045 (2001).

11. S. Allesina, A. Bodini, M. Pascual, Functional links and robustness in food webs. Philosophical Transactions of the Royal Society B: Biological Sciences 364, 1701–1709 (2009).

12. JoséM. Montoya, G. Woodward, M. C. Emmerson, R. V. Solé, Press perturbations and indirect effects in real food webs. Ecology 90, 2426–2433 (2009).

13. D. Sanders, E. Thébault, R. Kehoe, F. J. Frank van Veen, Trophic redundancy reduces vulnerability to extinction cascades. Proceedings of the National Academy of Sciences 115, 2419–2424 (2018).

14. A. Danet, M. Mouchet, W. Bonnaffé, E. Thébault, C. Fontaine, Species richness and food-web structure jointly drive community biomass and its temporal stability in fish communities. Ecology Letters 24, 2364–2377 (2021).

15. C. Damgaard, A Critique of the Space-for-Time Substitution Practice in Community Ecology. Trends in Ecology & Evolution 34, 416–421 (2019).

16. W. Bonnaffé, A. Danet, S. Legendre, E. Edeline, Comparison of size-structured and species-level trophic networks reveals antagonistic effects of temperature on vertical trophic diversity at the population and species level. Oikos, 1297–1309 (2021).

17. P. Chesson, J. J. Kuang, The interaction between predation and competition. Nature 456, 235–238 (2008).

18. M. I. O’Connor, et al., A general biodiversity–function relationship is mediated by trophic level. Oikos 126, 18–31 (2017).

19. J. Eschenbrenner, É. Thébault, Diversity, food web structure and the temporal stability of total plant and animal biomasses. Oikos 2023, e08769 (2023).

20. A. Danet, S. Kefi, T. F. Johnson, A. P. Beckerman, Response diversity is a major driver of temporal stability in complex food webs (2024) 10.1101/2024.08.29.610288 (September 2, 2024).

21. S. Wang, U. Brose, Biodiversity and ecosystem functioning in food webs: The vertical diversity hypothesis. Ecology Letters 21, 9–20 (2018).

22. A. Bodini, M. Bellingeri, S. Allesina, C. Bondavalli, Using food web dominator trees to catch secondary extinctions in action. Philosophical Transactions of the Royal Society B: Biological Sciences 364, 1725–1731 (2009).

23. M. R. Heithaus, A. Frid, A. J. Wirsing, B. Worm, Predicting ecological consequences of marine top predator declines. Trends in Ecology & Evolution 23, 202–210 (2008).

24. C. N. Johnson, J. L. Isaac, D. O. Fisher, Rarity of a top predator triggers continent-wide collapse of mammal prey: Dingoes and marsupials in Australia. Proceedings of the Royal Society B: Biological Sciences 274, 341–346 (2007).

25. J. A. Estes, et al., Trophic Downgrading of Planet Earth. Science 333, 301–306 (2011).

26. S. Wang, U. Brose, D. Gravel, Intraguild predation enhances biodiversity and functioning in complex food webs. Ecology 100, e02616 (2019).

27. S. Nie, et al., Will a large complex system be productive? Ecology Letters 26, 1325–1335 (2023).

28. S. Allesina, A. Bodini, Who dominates whom in the ecosystem? Energy flow bottlenecks and cascading extinctions. Journal of Theoretical Biology 230, 351–358 (2004).

29. R. L. Lindeman, The Trophic-Dynamic Aspect of Ecology. Ecology 23, 399–417 (1942).

30. D. Gravel, F. Massol, E. Canard, D. Mouillot, N. Mouquet, Trophic theory of island biogeography. Ecology Letters 14, 1010–1016 (2011).

31. J. A. Dunne, The network structure of food webs. Ecological networks: linking structure to dynamics in food webs, 27–86 (2006).

32. S. Yachi, M. Loreau, Biodiversity and ecosystem productivity in a fluctuating environment: The insurance hypothesis. Proceedings of the National Academy of Sciences 96, 1463–1468 (1999).

33. U. Brose, R. J. Williams, N. D. Martinez, Allometric scaling enhances stability in complex food webs. Ecology Letters 9, 1228–1236 (2006).

34. I. Lajaaiti, et al., EcologicalNetworksDynamics.jl: A Julia package to simulate the temporal dynamics of complex ecological networks. Methods in Ecology and Evolution n/a.

35. T. Woods, L. Comte, P. A. Tedesco, X. Giam, Testing the diversity–biomass relationship in riverine fish communities. Global Ecology and Biogeography 29, 1743–1757 (2020).

36. A. Hector, Plant Diversity and Productivity Experiments in European Grasslands. Science 286, 1123–1127 (1999).

37. D. Tilman, F. Isbell, J. M. Cowles, Biodiversity and Ecosystem Functioning. Annual Review of Ecology, Evolution, and Systematics 45, 471–493 (2014).

38. X. Wu, et al., The relationship between species richness and biomass changes from boreal to subtropical forests in China. Ecography 38, 602–613 (2015).

39. S. Li, et al., The relationship between species richness and aboveground biomass in a primary Pinus kesiya forest of Yunnan, southwestern China. PLOS ONE 13, e0191140 (2018).

40. J. B. Grace, et al., Integrative modelling reveals mechanisms linking productivity and plant species richness. Nature 529, 390–393 (2016).

41. D. M. Post, M. L. Pace, N. G. Hairston, Ecosystem size determines food-chain length in lakes. Nature 405, 1047–1049 (2000).

42. G. Takimoto, D. A. Spiller, D. M. Post, Ecosystem Size, but Not Disturbance, Determines Food-Chain Length on Islands of the Bahamas. Ecology 89, 3001–3007 (2008).

43. T. Poisot, N. Mouquet, D. Gravel, Trophic complementarity drives the biodiversity-ecosystem functioning relationship in food webs. Ecology Letters 16, 853–861 (2013).

44. M. Loreau, et al., Biodiversity as insurance: From concept to measurement and application. Biological Reviews 96, 2333–2354 (2021).

45. J. M. Montoya, S. L. Pimm, R. V. Solé, Ecological networks and their fragility. Nature 442, 259–264 (2006).

46. M. Barbier, M. Loreau, Pyramids and cascades: A synthesis of food chain functioning and stability. Ecology Letters 22, 405–419 (2019).

47. W. Bonnaffé, et al., The interaction between warming and enrichment accelerates food-web simplification in freshwater systems. Ecology Letters 27, e14480 (2024).

48. R. Froese, Cube law, condition factor and weight–length relationships: History, meta-analysis and recommendations. Journal of Applied Ichthyology 22, 241–253 (2006).

49. D. Gravel, T. Poisot, C. Albouy, L. Velez, D. Mouillot, Inferring food web structure from predator–prey body size relationships. Methods in Ecology and Evolution 4, 1083–1090 (2013).

50. U. Brose, et al., Predator traits determine food-web architecture across ecosystems. Nature Ecology & Evolution 3, 919–927 (2019).

51. S. Jennings, J. K. Pinnegar, N. V. C. Polunin, T. W. Boon, Weak cross-species relationships between body size and trophic level belie powerful size-based trophic structuring in fish communities. Journal of Animal Ecology 70, 934–944 (2001).

52. R. Froese, D. Pauly, et al., FishBase (Fisheries Centre, University of British Columbia, 2021).

53. G. G. Mittelbach, L. Persson, The ontogeny of piscivory and its ecological consequences. Canadian Journal of Fisheries and Aquatic Sciences 55, 1454–1465 (1998).

54. D. Claessen, C. V. Oss, A. M. de Roos, L. Persson, The Impact of Size-Dependent Predation on Population Dynamics and Individual Life History. Ecology 83, 1660–1675 (2002).

55. P. J. Hart, J. D. Reynolds, Handbook of Fish Biology and Fisheries: Fisheries (John Wiley & Sons, 2008).

56. J. D. Allan, M. M. Castillo, Stream ecology: Structure and function of running waters (Springer Science & Business Media, 2007).

57. E. Delmas, et al., Analysing ecological networks of species interactions: Analyzing ecological networks. Biological Reviews 94, 16–36 (2019).

58. R. A. Herendeen, “Ecological Network Analysis, Energy Analysis” in Encyclopedia of Ecology, S. E. Jørgensen, B. D. Fath, Eds. (Academic Press, 2008), pp. 1072–1083.

59. H. Rue, et al., Bayesian computing with INLA: A review. Annual Reviews of Statistics and Its Applications 4, 395–421 (2017).

60. S. Nakagawa, H. Schielzeth, A general and simple method for obtaining R2 from generalized linear mixed-effects models. Methods in Ecology and Evolution 4, 133–142 (2013).

61. D. M. LaHuis, M. J. Hartman, S. Hakoyama, P. C. Clark, Explained Variance Measures for Multilevel Models. Organizational Research Methods 17, 433–451 (2014).

62. A. Gelman, B. Goodrich, J. Gabry, A. Vehtari, R-squared for Bayesian Regression Models. The American Statistician 73, 307–309 (2019).

63. J. S. Lefcheck, piecewiseSEM: Piecewise structural equation modelling in r for ecology, evolution, and systematics. Methods in Ecology and Evolution 7, 573–579 (2016).

64. M. Murphy, semEff: Automatic Calculation of Effects for Piecewise Structural Equation Models (2021).

65. R. J. Williams, N. D. Martinez, Simple rules yield complex food webs. Nature 404, 180–183 (2000).

66. A. r. Ives, J. l. Klug, K. Gross, Stability and species richness in complex communities. Ecology Letters 3, 399–411 (2000).

67. M. Loreau, C. de Mazancourt, Species Synchrony and Its Drivers: Neutral and Nonneutral Community Dynamics in Fluctuating Environments. The American Naturalist 172, E48–E66 (2008).

