## Supplementary material for "Biodiversity temporal trends are reshaping food web structure and redundancy in riverine ecosystems"

#### 1 Disentangling functional and redundant trophic pathways

Species losses can trigger cascading extinctions by disrupting trophic interactions (1). To assess ecological network robustness, researchers often examine food web connectance — the proportion of possible trophic links that are realized. While higher connectance generally increases stability by providing multiple pathways for biomass flow, not all links contribute equally to robustness (2).

Trophic links can be categorized into two types based on their role in maintaining food web integrity. Functional links are those essential for connecting all species to basal resources — their removal directly triggers secondary extinctions. In contrast, redundant links provide alternative pathways but are not strictly necessary for maintaining basic connectivity. While the loss of redundant links doesn't cause immediate extinctions, it reduces the network's resilience to future perturbations (3). This distinction makes quantifying redundancy crucial for predicting ecosystem vulnerability.

To identify functional and redundant links, we employed dominator trees (4), a graph-theoretic approach that maps the minimal set of connections required to link all consumers to basal resources. Though once considered a niche tool, dominator tree algorithms are now readily accessible through common packages like R's `igraph::dominator_tree()`. Below, we demonstrate this method by replicating classic examples from the literature.

For implementation, we used a custom `compute_redundancy` function to quantify functional and redundant links. As an example, we applied this to the food-web from Bodini et al. (2009, see Appendix 1 therein), showing how easily these metrics can be derived from network data.

```r
compute_redundancy
#> function(rooted_graph, root = "R") {
#>
#>   # Compute the dominator tree
#>   dtree <- igraph::dominator_tree(rooted_graph, root = root, mode = "out")
#>
#>   # Compute number of links
#>   functional_links <- length(igraph::E(dtree$domtree))
#>   total_links <- length(igraph::E(rooted_graph))
#>   redundant_links <- total_links - functional_links
#>
#>   list(
#>       functional_links = functional_links,
#>       total_links = total_links,
#>       redundant_links = redundant_links,
#>       prop_redundant_links = redundant_links / total_links,
#>       prop_functional_links = functional_links / total_links
#>   )
#> }
```

```r
# Get the food-web example from Bodini et al. (2009), Appendix 1
g <- get_bodini_graph_appendix()
```

```r
# compute reduncancy
out <- compute_redundancy(g, root = "R")
as.data.frame(out)
#>   functional_links total_links redundant_links prop_redundant_links prop_functional_links
#> 1                4           7               3            0.4285714             0.5714286
```

Figure S1 provides visual comparisons between complete food-webs (left panels) and their functional-link subsets (right panels), reproducing key results from three seminal studies (3, 4). The accompanying Table S1 summarizes the proportion of redundant links in these systems, highlighting how dominator trees can simplify robustness assessments.

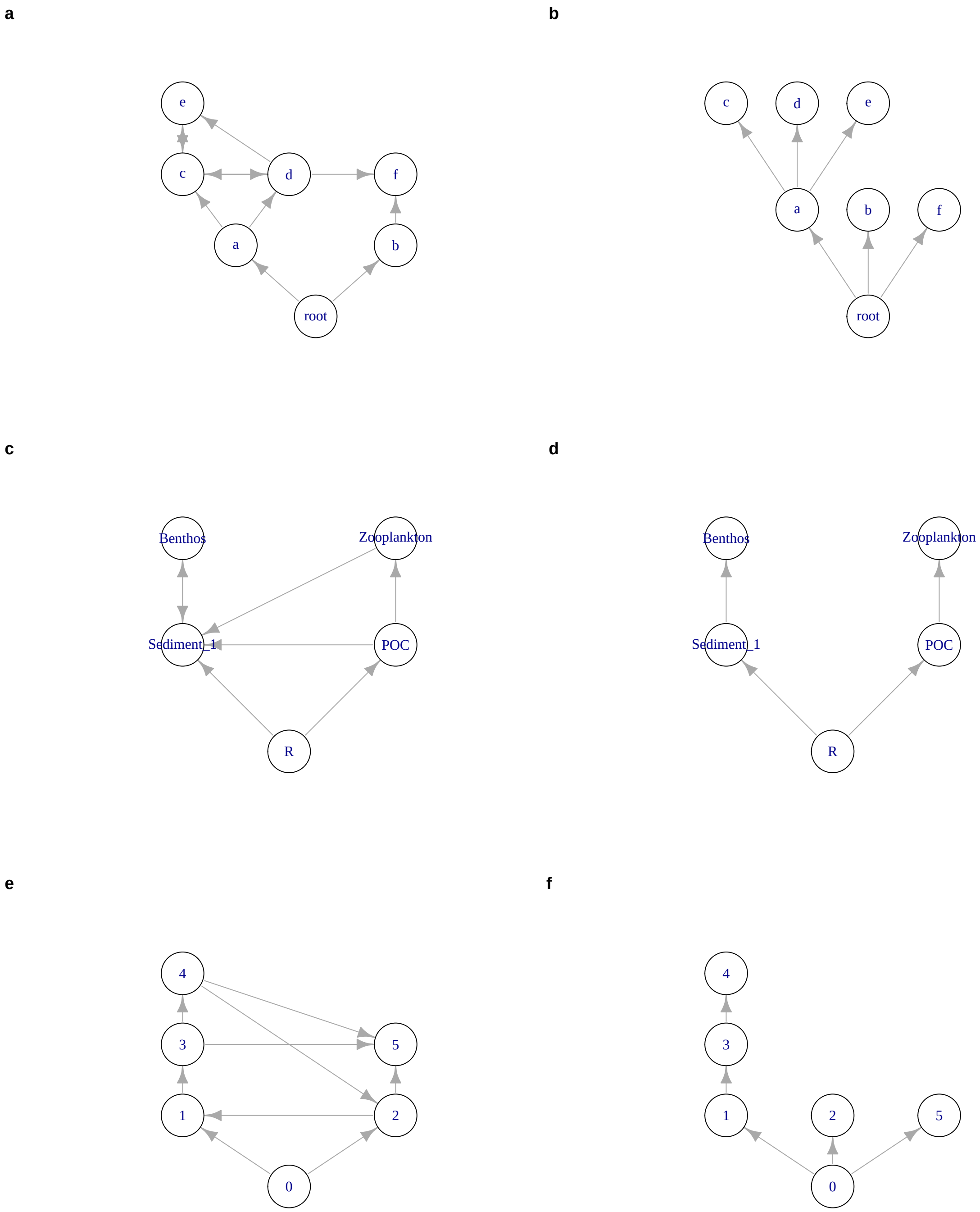


Figure S1: **Functional link identification using dominator trees**. Left panels show complete food-webs, while right panels display only functional links. (a,b) reproduce (4) Figure 1; (c,d) recreate their Appendix example; (e,f) replicate (4) Figure 1 (with minor differences noted in the original text, they could have removed further redundant links, i.e. $2 \rightarrow 5$, or $3 \rightarrow 5$).

### References


1. J. A. Dunne, R. J. Williams, N. D. Martinez, Network structure and biodiversity loss in food webs: Robustness increases with connectance. *Ecology Letters* **5**, 558–567 (2002).

Table S1: Computation of the number (Nb) of functional, redundant links, and their proportion (Prop.) from previous published food-webs.

| References | Nb links | Nb functional links | Nb redundant links | Prop. functional links | Prop. redundant links |
|---|---|---|---|---|---|
| Bodini et al. (2009) Supp material 1 | 7 | 4 | 3 | 0.57 | 0.43 |
| Bodini et al. (2009) Fig. 1 | 11 | 6 | 5 | 0.55 | 0.45 |
| Allesina et al. (2009) Fig. 1 | 9 | 5 | 4 | 0.56 | 0.44 |

2. S. Allesina, A. Bodini, Who dominates whom in the ecosystem? Energy flow bottlenecks and cascading extinctions. *Journal of Theoretical Biology* **230**, 351–358 (2004).

3. S. Allesina, A. Bodini, M. Pascual, Functional links and robustness in food webs. *Philosophical Transactions of the Royal Society B: Biological Sciences* **364**, 1701–1709 (2009).

4. A. Bodini, M. Bellingeri, S. Allesina, C. Bondavalli, Using food web dominator trees to catch secondary extinctions in action. *Philosophical Transactions of the Royal Society B: Biological Sciences* **364**, 1725–1731 (2009).

#### 2 Supplementary results: comparison of analysis in space and time

We further assessed the consistency of the reported associations among temporal trends by implementing two additional analyses. We first tested if the associations among the temporal trends within sites was holding across sites, i.e. using spatial variations in food web structure rather than temporal changes over time (space-for-time hypothesis). Second, we performed theoretical simulations with a bioenergetic food web model to cross-reference our site and species specific empirical data on riverine fish with a more generalized representation of food web dynamics known to represent accurate relationships among our focal variables. With those additional empirical and theoretical data, we assessed additional SEM models with the same structure as in Section 4.2 (see main text, Fig. 2a).

As in the temporal analysis, species richness had a positive effect on community biomass both in empirical data and in theoretical simulations (resp. $r_\delta$ = 0.47 [0.35,0.60] and 0.50 [0.46,0.53], Fig. 3a and S3). In both empirical data and theoretical simulations, community biomass had a positive effect on average trophic level (resp. $r_\delta$ = 0.35 [0.23,0.45] and 0.72 [0.69,0.75]), and redundancy (resp. $r_\delta$ = 0.32 [0.16,0.46] and 0.25 [0.21,0.30]), as in temporal analysis. We found weak evidence of a positive effect of spatial variation in community biomass on connectance in empirical data ($r_\delta$ = 0.11 [-0.04, 0.29]) but strong evidence in theoretical simulations ($r_\delta$ = 0.26 [0.23,0.30], Fig. 3a).

The effects of spatial variations in species richness on food web structure mostly aligned with those from the temporal trend analysis, except for average trophic level. We found a direct negative effect of species richness on connectance both using data and model simulations (resp. $r_\delta$ = -0.60 [-0.65,-0.52] and -0.77 [-0.79,-0.75], Fig. 3a), on average trophic level (resp. $r_\delta$ = -0.32 [-0.54,-0.18] and -0.21 [-0.26,-0.18]), but a direct positive effect on redundancy (resp. $r_\delta$ = 0.38 [0.19,0.49] and 0.38 [0.32,0.43]). Species richness had an indirect positive effect of species richness through community biomass, which added up to strong total negative effects on connectance in empirical data and simulations (resp. $r_\delta$ = -0.55 [-0.59,-0.43] and -0.64 [-0.66,-0.61]) and positive on redundancy (resp. $r_\delta$ = 0.53 [0.32,0.62] and 0.50 [0.47,0.54]). We found that species richness had a total positive effect on average trophic level in bioenergetic model, in agreement with temporal trends (resp. $r_\delta$ = 0.15 [0.10,0.19]), but a total negative effect in spatial variations (resp. $r_\delta$ = -0.16 [-0.38,-0.04]).

### Supplementary figures

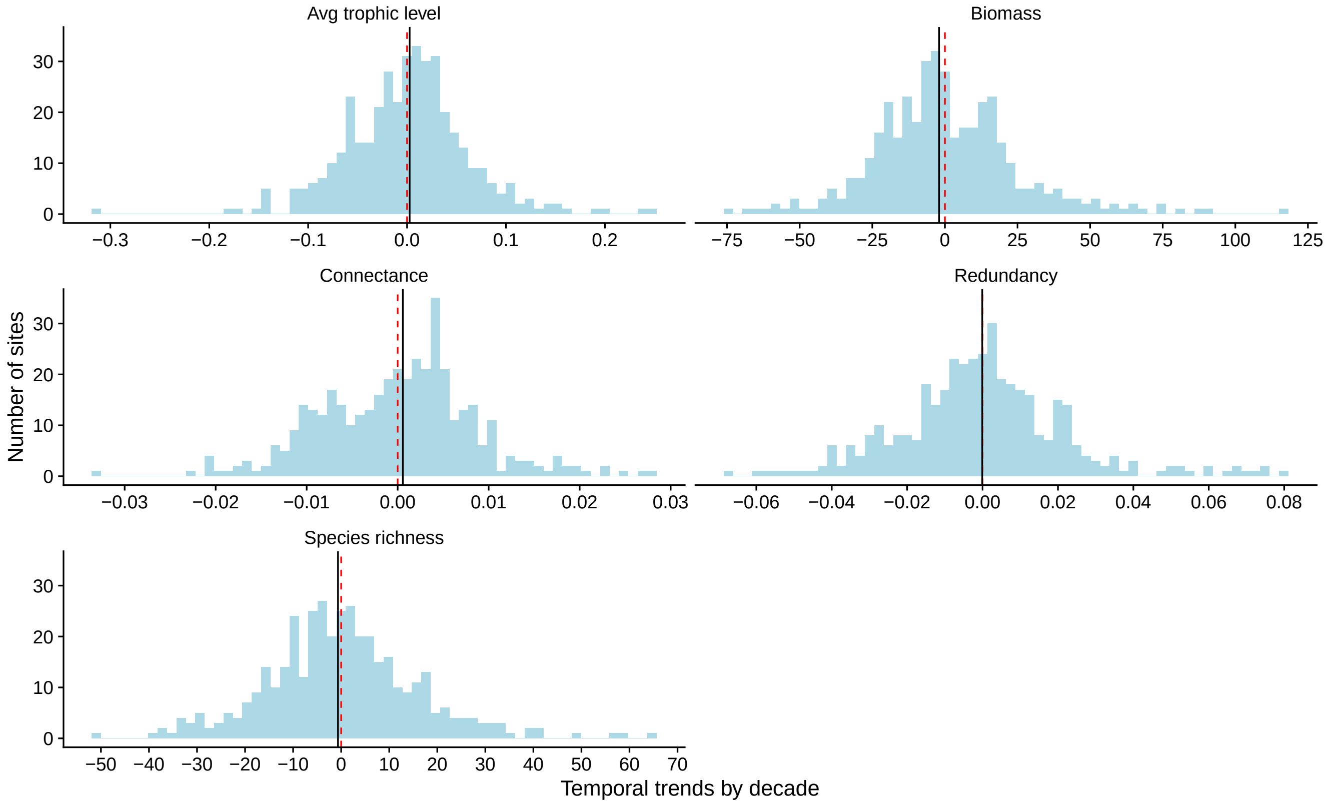


Figure S2: Distribution of temporal trends per decade of the food-web structure by site estimated from the Bayesian models using Best Linear Unbiased Prediction (BLUP). Species richness and biomass temporal trends are reported in percentage change per decade. Dashed red line and solid black line respectively indicate 0 and median change.

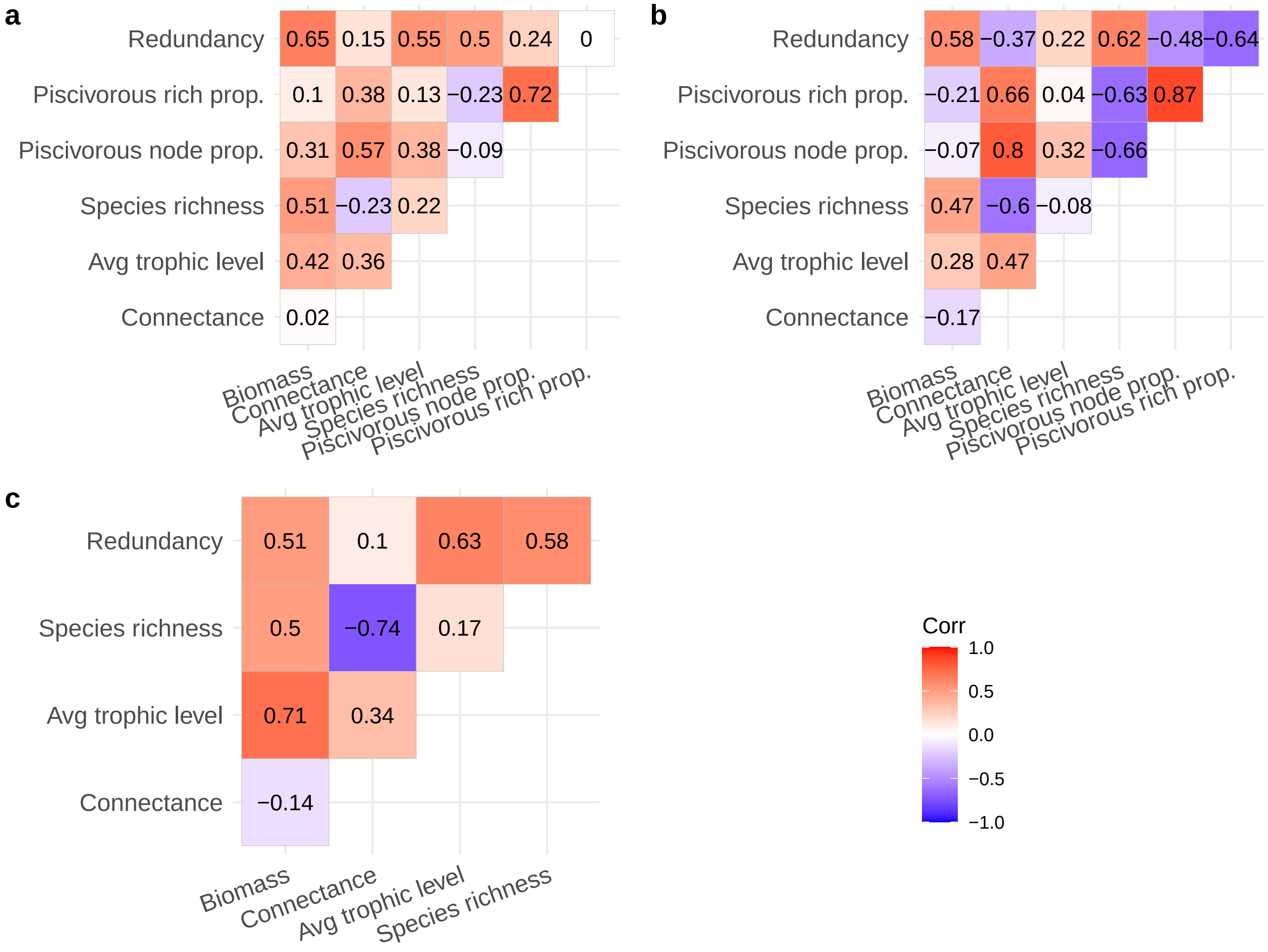


Figure S3: Correlation among food-web metrics for a) temporal trends by site, b) median values by site (cf Main Methods), c) bioenergetic model. Pisc: piscivorous, rich.: species richness, node: node of the food-web corresponding to a trophic species, prop.: proportion.

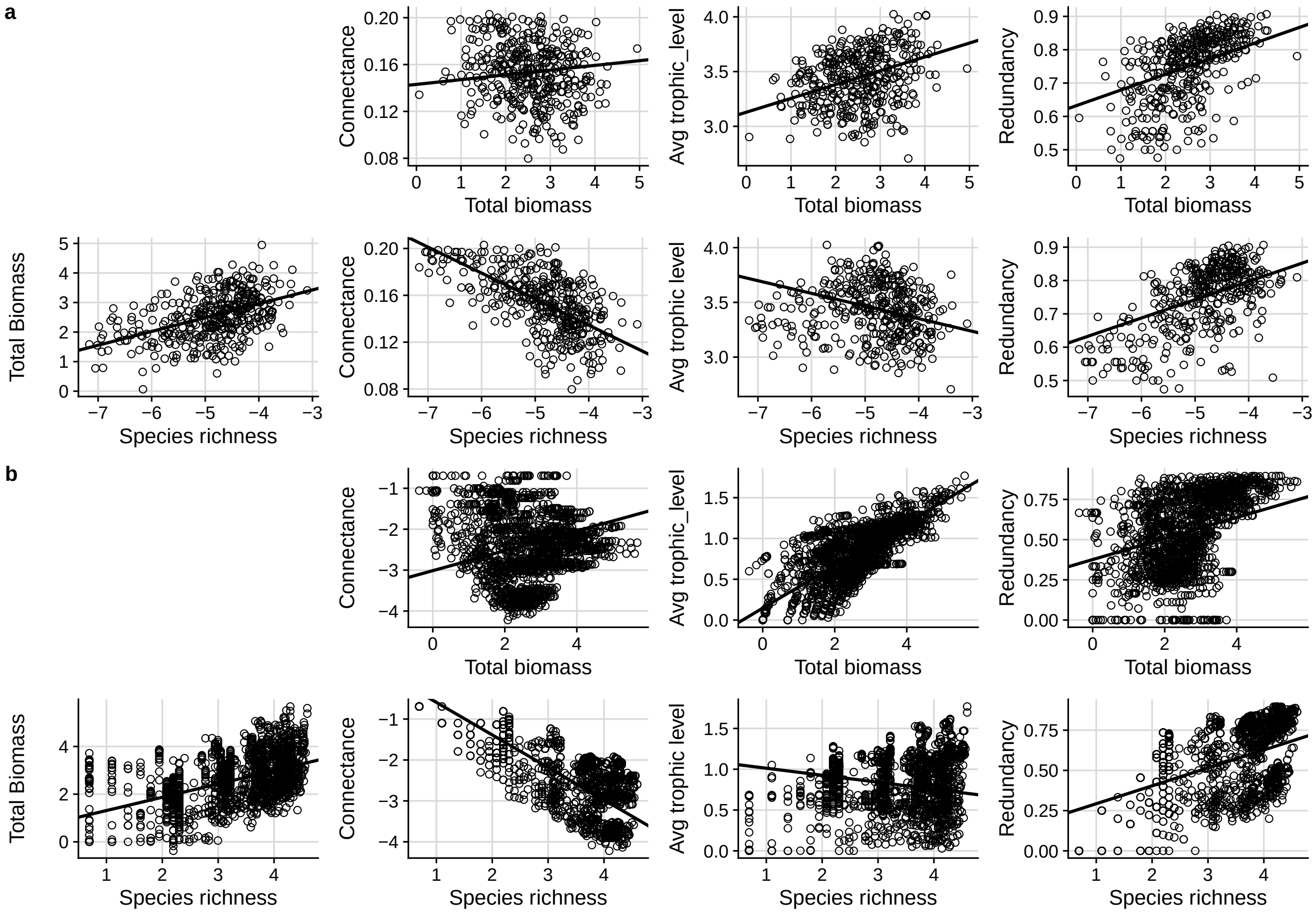


Figure S4: Direct effects of biomass and species richness on food-web structure, derived from the structural equation models for the (a) spatial and (b) bioenergetic model. Points display the data, black lines display the predicted relationships.

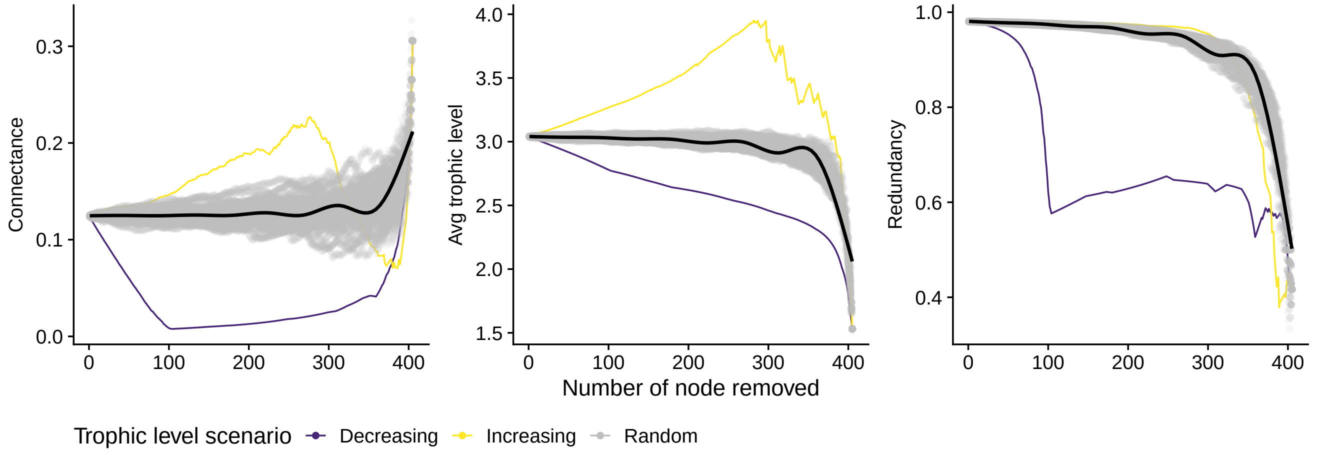


Figure S5: Simulations of trophic species extinctions in the metaweb according to three scenarios: random extinction sequence (grey dots), extinction sequence by decreasing order of trophic level, extinction sequence by increasing order of trophic level. The black line represents the average prediction of a GAM linear model for the 50 random extinction sequences. Please note that the results are qualitatively similar that when considering species (see Fig. 3b, main text) instead of trophic species.

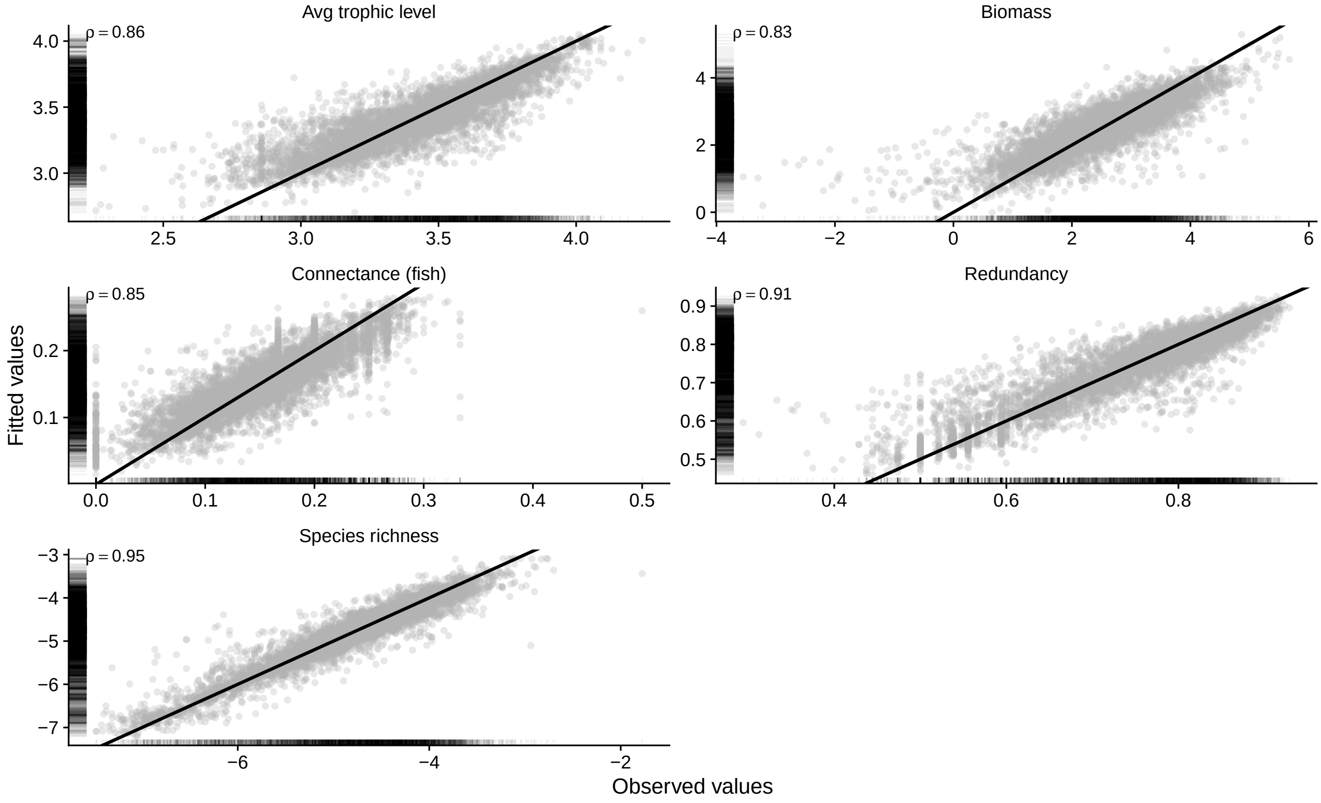


Figure S6: Fitted versus observed values from the linear models estimating the temporal trends of the food-web structure. The black line depicts the first bissector. Pearson's correlation coefficients are displayed.

#### Supplementary tables

Table S2: Summary distribution of community metrics and fish monitoring variables. Q1 and Q3 are the first and third quartiles of the distributions.

| Variable | Median (Q1, Q3) | (Min, Max) |
|---|---|---|
| **Community metrics** | | |
| Connectance | 0.16 ( 0.13, 0.18) | (0.05, 0.25) |
| Piscivorous node prop. | 0.27 ( 0.20, 0.36) | (0.00, 0.83) |
| Piscivorous rich prop. | 0.33 ( 0.25, 0.43) | (0.00, 1.00) |
| Redundancy | 0.77 ( 0.67, 0.83) | (0.29, 0.93) |
| Avg trophic level | 3.44 ( 3.22, 3.64) | (2.26, 4.24) |
| Species richness | 7.00 ( 4.00, 11.00) | (1.00, 25.00) |
| Biomass (kg) | 9.35 ( 4.43, 18.92) | (0.00, 182.32) |
| Nb of nodes | 30.00 ( 20.00, 41.00) | (8.00, 92.00) |
| **Summary of fish monitoring** | | |
| Nb of individuals | 343.00 (174.00,640.00) | (1.00,9961.00) |
| Baseline year | 1995.00 (1995.00,1998.50) | (1995.00,2009.00) |
| Completeness | 0.92 ( 0.79, 1.00) | ( 0.22, 1.00) |
| Sampling number | 12.00 ( 11.00, 18.00) | ( 1.00, 24.00) |
| Year span | 14.00 ( 12.00, 23.00) | ( 1.00, 24.00) |

Table S3: Marginal and Conditional R-squared for the INLA models estimating the temporal trends of the food-web structure. Mean [95% Credible Interval].

| Response | Marg. Rsq | Cond. Rsq |
|---|---|---|
| Biomass | 0.03 [0.01,0.06] | 0.67 [0.66,0.68] |
| Connectance | 0.03 [0.01,0.06] | 0.70 [0.69,0.71] |
| Avg trophic level | 0.02 [0.01,0.04] | 0.77 [0.76,0.77] |
| Species richness | 0.01 [0.00,0.02] | 0.89 [0.89,0.89] |
| Redundancy | 0.02 [0.01,0.04] | 0.79 [0.79,0.80] |

Table S4: Direct and total Standardized effects derived from the structural equation models. [95% Confidence Interval]. The confidence intervals were obtained with bootstrap (see Methods).

| Response | Predictor | Direct | Total |
|---|---|---|---|
| **Temporal** | | | |
| Biomass | Species richness | 0.53 [0.42; 0.61] | 0.53 [0.42; 0.61] |
| Connectance | Species richness | -0.28 [-0.39; -0.16] | -0.2 [-0.28; -0.1] |
| | Biomass | 0.17 [0.04; 0.28] | 0.17 [0.04; 0.28] |
| Avg trophic level | Species richness | 0.02 [-0.1; 0.11] | 0.21 [0.08; 0.31] |
| | Biomass | 0.37 [0.27; 0.46] | 0.37 [0.27; 0.46] |
| Redundancy | Species richness | 0.19 [0.08; 0.29] | 0.43 [0.34; 0.5] |
| | Biomass | 0.46 [0.37; 0.54] | 0.46 [0.37; 0.54] |
| **Spatial** | | | |
| Biomass | Species richness | 0.47 [0.35; 0.6] | 0.47 [0.35; 0.6] |
| Connectance | Species richness | -0.6 [-0.65; -0.52] | -0.55 [-0.59; -0.43] |
| | Biomass | 0.11 [-0.04; 0.29] | 0.11 [-0.04; 0.29] |
| Avg trophic level | Species richness | -0.32 [-0.54; -0.18] | -0.16 [-0.38; -0.04] |
| | Biomass | 0.35 [0.23; 0.45] | 0.35 [0.23; 0.45] |
| Redundancy | Species richness | 0.38 [0.19; 0.49] | 0.53 [0.32; 0.62] |
| | Biomass | 0.32 [0.16; 0.46] | 0.32 [0.16; 0.46] |
| **Theoretical model** | | | |
| Biomass | Species richness | 0.5 [0.46; 0.53] | 0.5 [0.46; 0.53] |
| Connectance | Species richness | -0.77 [-0.79; -0.75] | -0.64 [-0.66; -0.61] |
| | Biomass | 0.26 [0.23; 0.3] | 0.26 [0.23; 0.3] |
| Avg trophic level | Species richness | -0.21 [-0.26; -0.18] | 0.15 [0.1; 0.19] |
| | Biomass | 0.72 [0.69; 0.75] | 0.72 [0.69; 0.75] |
| Redundancy | Species richness | 0.38 [0.32; 0.43] | 0.5 [0.47; 0.54] |
| | Biomass | 0.25 [0.21; 0.3] | 0.25 [0.21; 0.3] |

Table S5: Marginal and conditional R squared for the Structural Equation Models.

| Type | Response | Marg. R2 | Cond. R2 |
|---|---|---|---|
| Temporal | Biomass | 0.28 | NA |
| | Connectance | 0.08 | NA |
| | Avg trophic level | 0.20 | NA |
| | Redundancy | 0.47 | NA |
| Spatial | Biomass | 0.22 | 0.26 |
| | Connectance | 0.38 | 0.39 |
| | Avg trophic level | 0.14 | 0.28 |
| | Redundancy | 0.46 | 0.48 |
| Theoretical model | Biomass | 0.25 | NA |
| | Connectance | 0.61 | NA |
| | Avg trophic level | 0.54 | NA |
| | Redundancy | 0.40 | NA |

Table S6: Standard deviation associated to the random effects and the residual error of the models. Mean [95% Credible Interval].

| Response | Term | S.D. |
|---|---|---|
| Biomass | Time (site nested in basin) | 0.034 [0.03,0.04] |
| | Time (basin) | 0.008 [0.005,0.03] |
| | Intercept (site nested in basin) | 0.773 [0.72,0.839] |
| | Intercept (basin) | 0.222 [0.132,1.15] |
| | Error | 0.58 [0.569,0.592] |
| Connectance | Time (site nested in basin) | 0.001 [0.001,0.001] |
| | Time (basin) | 0.004 [0.003,0.012] |
| | Intercept (site nested in basin) | 0.024 [0.022,0.026] |
| | Intercept (basin) | 0.006 [0.004,0.014] |
| | Error | 0.017 [0.016,0.017] |
| Avg trophic level | Time (site nested in basin) | 0.009 [0.007,0.01] |
| | Time (basin) | 0 [NA,NA] |
| | Intercept (site nested in basin) | 0.237 [0.221,0.258] |
| | Intercept (basin) | 0.133 [0.12,0.154] |
| | Error | 0.152 [0.15,0.156] |
| Species richness | Time (site nested in basin) | 0.02 [0.018,0.023] |
| | Time (basin) | 0.008 [0.005,0.026] |
| | Intercept (site nested in basin) | 0.668 [0.624,0.721] |
| | Intercept (basin) | 0.294 [0.199,0.724] |
| | Error | 0.255 [0.25,0.26] |
| Redundancy | Time (site nested in basin) | 0.003 [0.003,0.004] |
| | Time (basin) | 0.004 [0.003,0.058] |
| | Intercept (site nested in basin) | 0.09 [0.084,0.097] |
| | Intercept (basin) | 0.027 [0.019,0.056] |
| | Error | 0.048 [0.047,0.049] |

Table S7: Variance Inflation Factor for the more complex linear models included in the Structural Equation Models.

| Model | Biomass | Species richness |
|---|---|---|
| Temporal | 1.39 | 1.39 |
| Spatial | 1.25 | 1.25 |
| Theoretical model | 1.33 | 1.33 |
